# *Vibrio alginolyticus* is the pathogen of ‘Baotou’ disease causing serious damage to *Gracilariopsis lemaneiformis* cultivation in China

**DOI:** 10.1101/2024.08.27.609959

**Authors:** Tong Pang, Feng Wang, Qunqun Guo, Mengjie Zhang, Yuanyuan Sun, Jianguo Liu

**Author notes:** Tong Pang and Feng Wang contributed equally to this work. Author order was determined on the basis of seniority. Corresponding author. Tong Pang, Institute of Oceanology, Chinese Academy of Sciences, 7 Nanhai Road, Qingdao, China, 266071.; Jianguo Liu, Institute of Oceanology, Chinese Academy of Sciences, 7 Nanhai Road, Qingdao, China, 266071.

## Abstract

*Gracilariopsis lemaneiformis* and *Saccharina japonica*, the two very important cultivated economic seaweeds, are often co-cultured in China due to their growth periods overlapped for about 3 months. These two seaweeds are often plagued with bleaching diseases during their high-density commercial cultivation. In recent years, a disease called ‘Baotou’ has been causing large-scale yield reductions of *G. lemaneiformis* in China. Interestingly, *Vibrio alginolyticus* once reported to be beneficial to *S. japonica* by reducing the bleaching disease risk, was strongly proved to be the pathogen causing ‘Baotou’ disease in this study. Analysis of 16S rRNA gene profiling revealed that *V. alginolyticus* was the most abundant and dominant bacterium on the algal thalli suffering from ‘Baotou’ disease, whereas its presence was scarcely detected on healthy thalli. Scanning electron microscope (SEM) analysis revealed that a large number of *V. alginolyticus* cells were found to be attached to the algal thalli with ‘Baotou’ disease and the rotten thalli acquired by the lab infection treatment. *V. alginolyticus* could cause the rotten symptoms which were consistent with those of ‘Baotou’ disease. According to Koch’s postulates, *V. alginolyticus* was identified as the pathogen causing ‘Baotou’ disease.

**IMPORTANCE:** A highly contagious disease known as ‘Baotou’ disease has persistently triggered significant yield reductions in *G. lemaneiformis* throughout China. The pathogen of ‘Baotou’ disease was isolated and identified as *V. alginolyticus* which was once reported to be beneficial to *S. japonica* by reducing the bleaching disease risk. The study indicates that *V. alginolyticus* is closely involved in the competition or co-existence between *G. lemaneiformis* and *S. japonica* co-cultured in China, and provides a novel perspective and evidence that bacteria are closely involved in the competition or co-existence among different species of seaweeds that grow together.

## Introduction

Seaweeds can grow rapidly in the offshore farm without occupying land, and are rich in various chemical compositions that are useful to humans (1,2). For decades, its extensive application in various sectors, including food, feed, biofuels, chemicals, nutraceuticals, medicines, cosmetics, and environmental bioremediation, has promoted worldwide seaweed production (1,2). The productivity of seaweed increased significantly from 10.6 million tons in 2000 to 35 million tons in 2020, dominated especially by *Saccharina japonica*, *Eucheumatoid* species and *Gracilarioids* species (3). The *Gracilariopsis lemaneiformis*, a species of *Gracilarioids* algae, is widely distributed in coastal countries all over the world, such as the China, America. (4,5). It not only serves as a raw material for agar production and feed for abalone aquaculture but also plays a crucial role in regulating and restoring the ecological environment, thus bearing significant importance for human production and livelihoods (6,7). The total area of cultivated *G. lemaneiformis* in China exceeded 13,000 hectares in 2022 (8). *G. lemaneiformis* has become the second largest main seaweed species planted after *Eucheumatoid* species (9).

In recent years, with the expansion of *G. lemaneiformis* breeding industry, various diseases and pests have been frequently and continuously affecting the cultivation of *G. lemaneiformis.* Various diseases such as apical necrosis and branch whitening have led to a reduction in the production of *G. lemaneiformis* (10,11). Especially in 2023, the impact of a disease called ‘Baotou’ disease resulted in a 40% reduction in *G. lemaneiformis* production compared to that in 2022 (12). Besides, *Ceramium filrula,* a epiphytes alga that is often attached to *G. lemaneiformis*, which competes for nutrients and light from the host and slows down *G. lemaneiformis* growth (13). Herbivores, such as rabbitfish (*Siganus canaliculatus*), was also a challenge in *G. lemaneiformis* cultivation (14).

Intensive studies have been conducted on the pathogens of various algal rot diseases (15). Alginic acid decomposing bacteria could cause rot disease when *Saccharina japonica* sporeling was incubated in stressful environmental conditions (16). *Pseudoalteromonas, Vibrio* and *Halomonas* may be the potential pathogenic bacteria associated with the Hole-Rotten Disease of *S. japonica* (17). *V. alginolyticus* was found to be a beneficial bacterium to *S. japonica* and reduces the bleaching disease risk of *S. japonica* (18). *Vibrio* sp. P11 promoted ice-ice disease in *K. alvarezii* (19,20). Syafitri et al. (2017) found that the ice-ice disease of *K. alvarezii* was mainly caused by three types of bacteria: *Alteromonas macleodii, Pseudoalteromonas issachenkonii, and Aurantimonas coralicida*, among which *A. macleodii* exhibited the strongest pathogenicity (21). The yellow spot disease (YSD) that occurred in *conchocelis* sporeling cultures of *Pyropia* could be induced by *Vibrio medierranei* 117-T6 (22). Ding and Ma (2005) reported that in the red rot diseases of *Porphyra yezoensis*, both *Pythium porphyrae* and *Olpidiopsis* sp. were observed as dominant bacteria (23). Agar-digesting Vibrio species from the rotten thallus of *Gracilariopsis heteroclada* was isolated and characterized by Lavina-Pitogo (1992) and Martinez and Padilla (2016) (24,25). *Thalassospira* sp. and *Vibrio parahaemolyticus*, which could induce the necrosis of healthy tips on *G. lemaneiformis*, were isolated by Sun et al. (2011) (11). *Agarivorans albus*, *Aquimarina latercula (T)* and *Brachybacterium* sp., which can bleach the healthy *G. lemaneiformis*, were isolated from the bleached *G. lemaneiformis* (10).

Despite the ‘Baotou’ disease continuously and seriously affecting the cultivation of *G. lemaneiformis*, there is still a lack of research on the pathogen. Considering the enormous impact of ‘Baotou’ disease on the production of *G. lemaneiformis*, the pathogen of ‘Baotou’ disease was isolated and identified in this study.

## Results

### Bacterial communities on healthy and rotten algae

*Vibrio alginolyticus* is the dominant bacterium on the rotten samples (Fig 1 a) and accounted for 72.32%, 54.37% and 36.53% of the total microbial community on samples R1, R2 and R3 respectively (Fig 1a). In contrast, *V. alginolyticus* is not detected on the healthy samples of H1 and H3, and only accounted for only 0.56% of the H2 sample. On average, the *V. alginolyticus* accounted 54.41% of the total microbial community on rotten samples, which was significantly higher than that on the heathy samples (0.189%, Fig 1b).

**Fig. 1.**
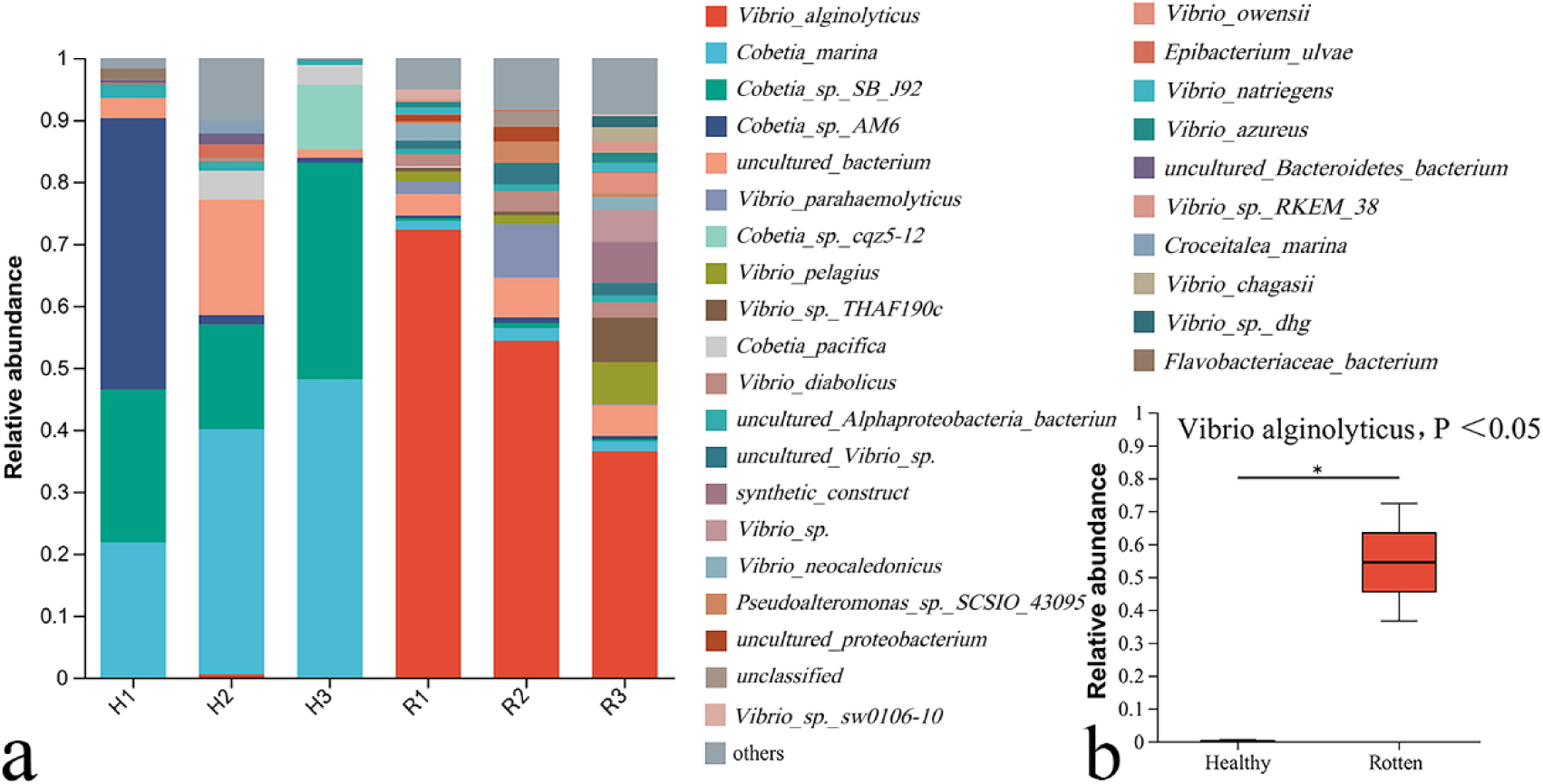
Bacterial communities on healthy and rotten *G. lemaneiformis* thalli. (a) The relative abundance of detectable bacteria on healthy (H) and rotten (R) thalli, (b). The relative abundance of *V. alginolyticus* on healthy and rotten thalli.

### Bacterial communities in seawater near healthy and rotten algae

*V. alginolyticus* accounted for 5.58%, 11.78% and 4.96% of the total microbial community on samples RS1, RS2 and RS3 respectively (Fig 2a). In contrast, *V. alginolyticus* is not detected on the healthy samples of HS2 and HS3, and only accounted for 0.53% of the HS1 sample. On average, the *V. alginolyticus* accounted 7.5% of the total microbial community in the seawater near rotten samples, which was significantly higher than that in the seawater near heathy samples (0.18%, Fig 2b).

**Fig. 2.**
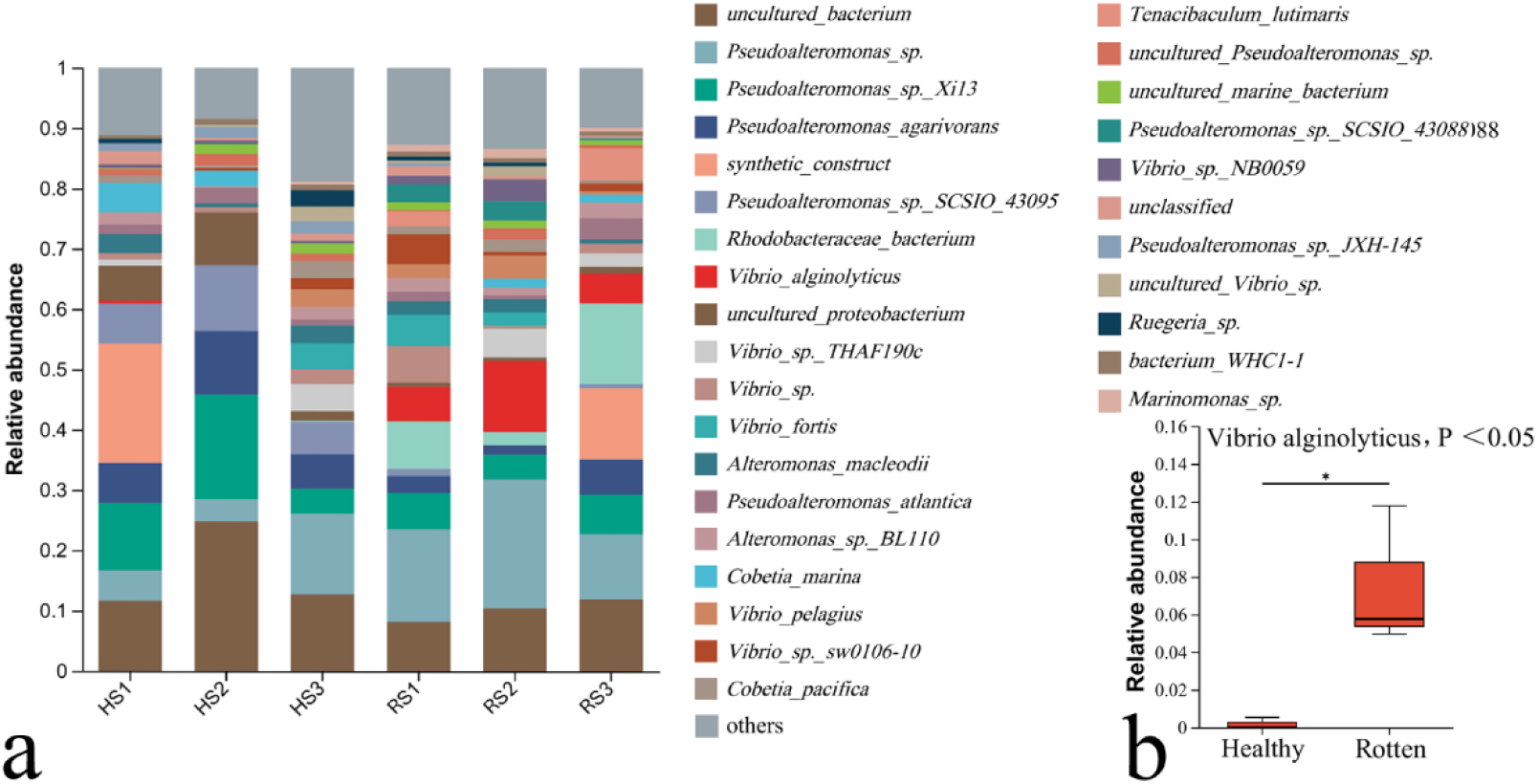
Bacterial communities in seawater near healthy (HS) and rotten (RS) *G. lemaneiformis* thalli. (a) The relative abundance of detectable bacteria in the seawater bacterial community near healthy and rotten thalli, (b). The relative abundance of V. alginolyticus in the seawater near healthy and rotten thalli.

### Isolated bacteria and their pathogenicity

49 colonies were obtained for pathogenicity testing, of which 10 strains could induce the rotten of *G. lemaneiformis*. ZB10 exhibited the highest pathogenicity in the 10 strains of bacteria. Six different species of bacteria, which could cause the rotting of *G. lemaneiformis* thalli, were identified by the 16S rRNA phylogenetic analysis (Table 1, supplementary Figs). The pathogenicity of ZB10 and ZB7 was significantly higher than that of the other species at 72 h. The main rotting part caused by ‘Baotou’ disease is the trunk of the algae (Fig. 3a, b). The symptom caused by the ZB10 and ZB7 was the rotting of the main trunk of the algae, that was consistent with the ‘Baotou’ disease occurring in farms (Fig. 3a, b).

**Table 1.**
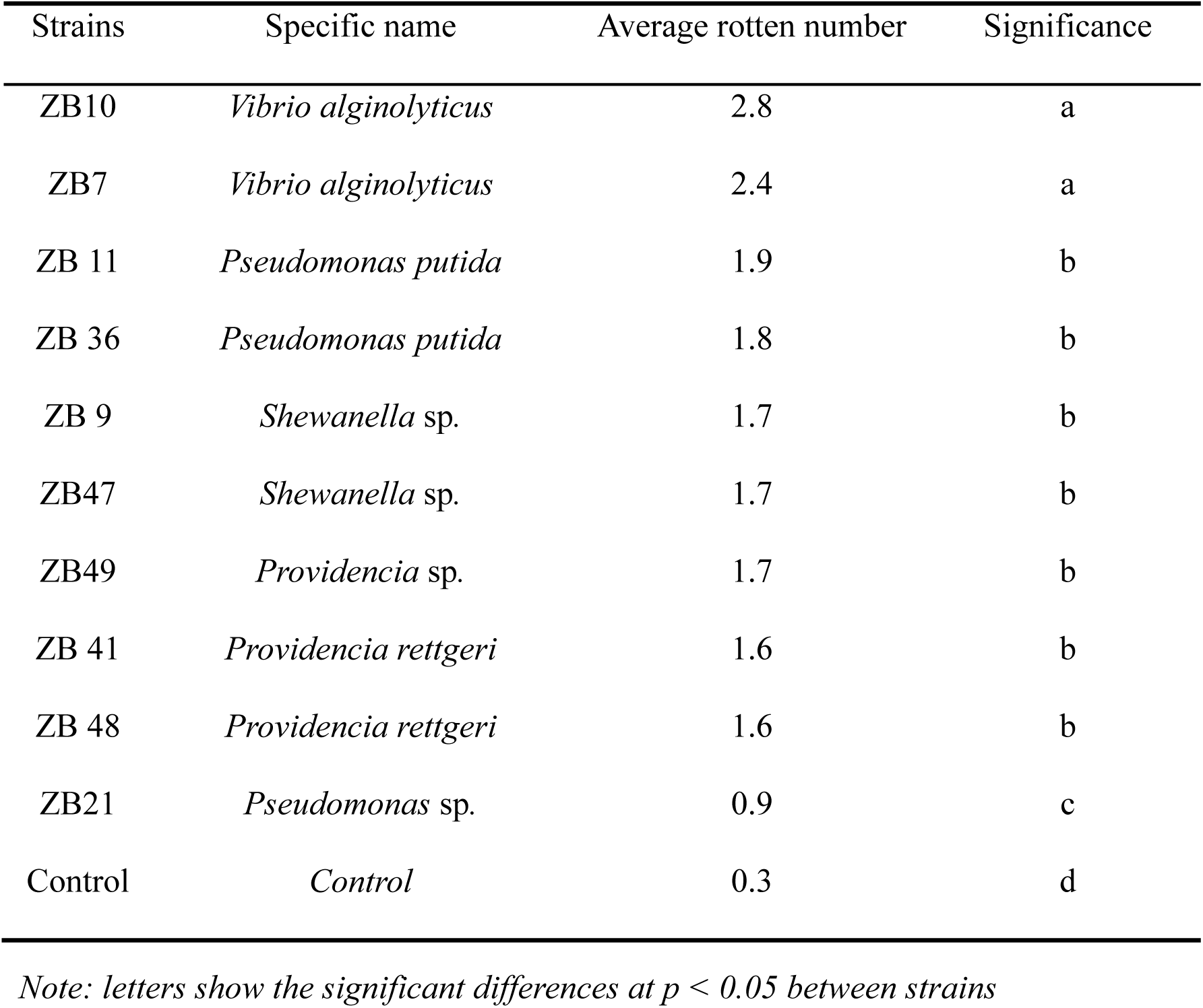
Isolated pathogenic bacteria and their pathogenicity at 72 hours. (n=9)

| Strains | Specific name | Average rotten number | Significance |
| --- | --- | --- | --- |
| ZB10 | *Vibrio alginolyticus* | 2.8 | a |
| ZB7 | *Vibrio alginolyticus* | 2.4 | a |
| ZB 11 | *Pseudomonas putida* | 1.9 | b |
| ZB 36 | *Pseudomonas putida* | 1.8 | b |
| ZB 9 | *Shewanella* sp. | 1.7 | b |
| ZB47 | *Shewanella* sp. | 1.7 | b |
| ZB49 | *Providencia* sp. | 1.7 | b |
| ZB 41 | *Providencia rettgeri* | 1.6 | b |
| ZB 48 | *Providencia rettgeri* | 1.6 | b |
| ZB21 | *Pseudomonas* sp. | 0.9 | c |
| Control | *Control* | 0.3 | d |

Note: letters show the significant differences at $p < 0.05$ between strains

**Fig. 3.**
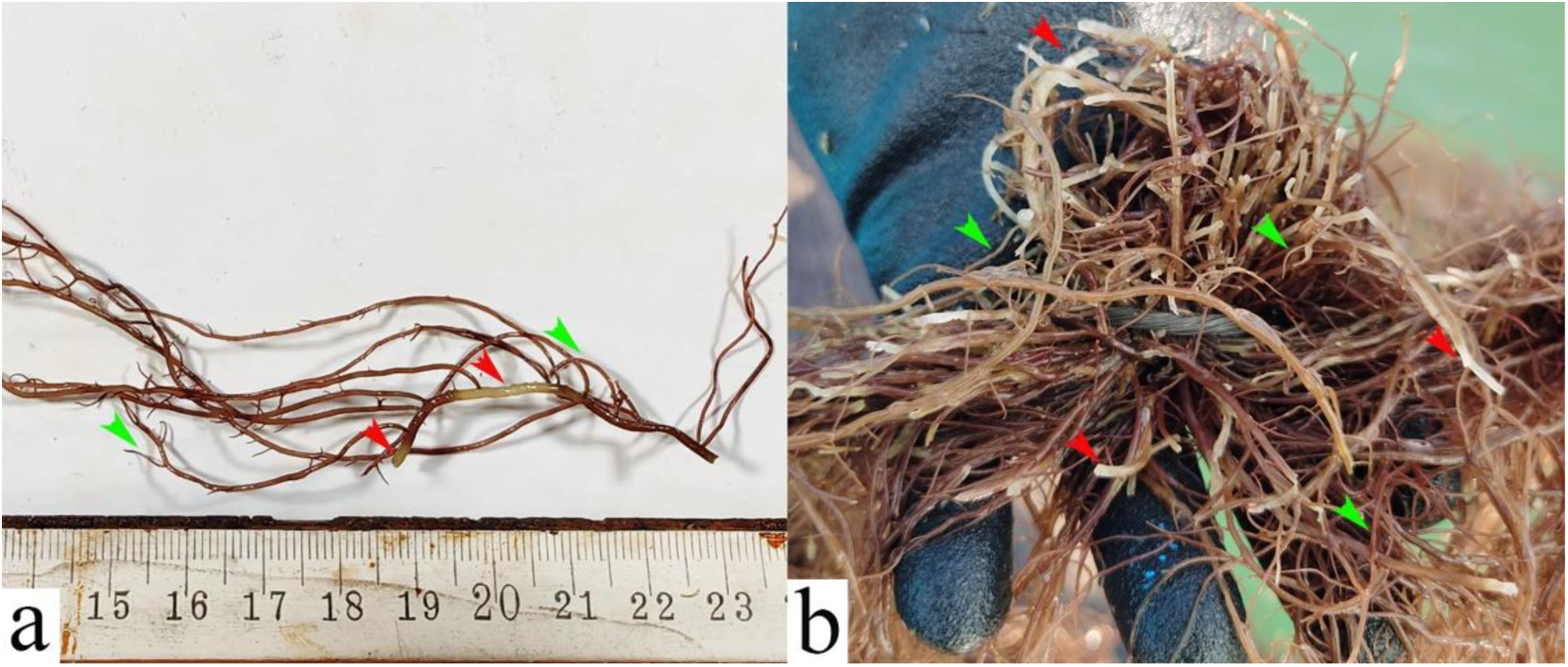
Rotten *G. lemaneiformis* after infection. (a) Rotten *G. lemaneiformis* infected by strain ZB10 in the lab. (b) Rotten *G. lemaneiformis* with ‘Baotou’ disease in the farm. Red arrows point to the rotten main trunk, blue arrows point to healthy tips.

### Morphology of the pathogenic bacteria

The colonies of ZB10 were bright yellow protruding on TCBS medium, with a single colony diameter of about 2-3 mm and growing diffusely (Fig 4 a). The cell shapes of ZB10 were rod-shaped or spherical (Fig 4 b, c d). One curved flagellum was found on the ZB10 under the TEM. The cell sizes of *V. alginolyticus* varied greatly, with large ones reaching 4 μm long and small ones only 1 μm long or even less. (Fig. 4 c and d). All the above morphological characteristics were consistent with the morphological characteristics of *V. alginolyticus* reported by Zhao et al (30). Combined the 16S rRNA phylogenetic analysis and morphological results, ZB10 was identified as *V. alginolyticus*. A large number of *V. alginolyticus* cells were found to be attached to the decaying thalli surface of *G. lemaneiformis* after 72 h of co-culturing and infecting with the *V. alginolyticus* (Fig 4 e). In addition, *V. alginolyticus* can be easily re-isolated from the decayed algal thalli infected in the laboratory. Furthermore, a large number of rod-shaped and spherical bacterial cells similar to those found in the lab infection experiments were also discovered on the *G. lemaneiformis* thalli with ‘Baotou’ disease (Fig. 4 f). However, the size of the *V. alginolyticus* found on the ‘Baotou’ disease algae collected from the farm was smaller than that induced in the lab (Fig. 4 e, f).

**Fig. 4.**
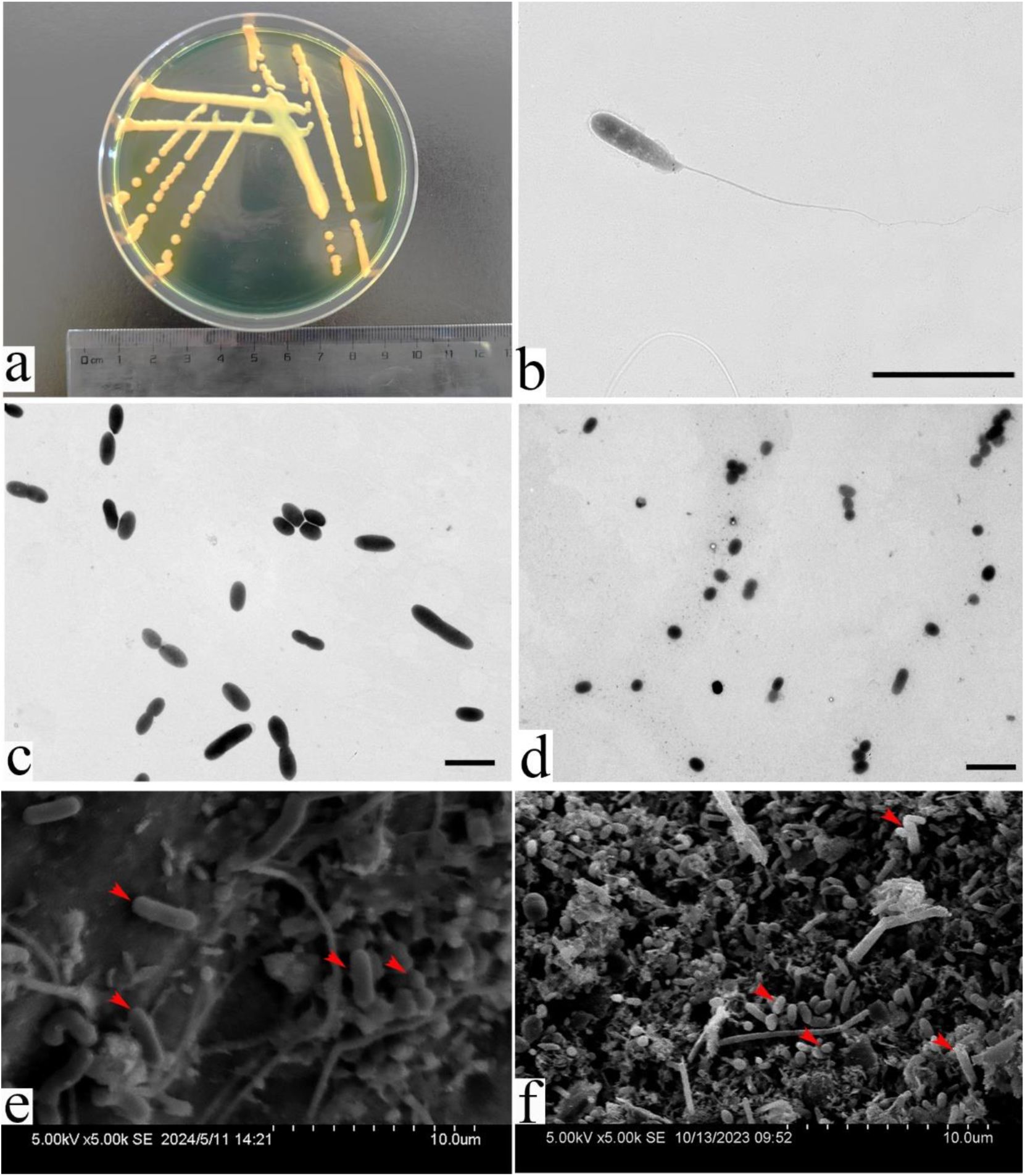
Morphology of the bacteria ZB10. (a) Colony morphology of ZB10 on TCBS medium. (b) Morphology of the bacteria ZB10 under the TEM with a magnification of 4K. (c) Large cells of ZB10 under the TEM with a magnification of 0.7K. (d) Small cells of ZB10 under the TEM with a magnification of 0.7K, small ones are only 1 μm long. (e) Surface views of the rotten algal thalli derived from lab infection treatment under SEM with a magnification of 5K. (f) Surface views of the algal thalli from farm with ‘Baotou’ disease under SEM with a magnification of 5K. Bars: b c and d :5μm. Red arrows point to pathogenic bacteria *V. alginolyticus*.

## Discussion

Due to the ‘Baotou’ disease, Chinese *G. lemaneiformis* yield decreased by at least 40% in 2023 compared with that in 2022 (12). Two strains of pathogenic bacteria, *Thalassospira* sp. and *Vibrio parahaemolyticus*, which could cause the tip rotting of *G. lemaneiformis* was isolated by Sun et al (11). Pathogenic bacteria, *Agarivorans albus*, *Aquimarina latercula* and *Brachybacterium* sp., which could induce the bleaching of *G. lemaneiformis*, were isolated by Liu et al (10). However, the pathogenic bacteria isolated by Sun et al (11) and Liu et al (10) were not the dominant bacteria on the rotten *G. lemaneiformis* with ‘Baotou’ disease (Fig. 1a), and were also not found in the seawater of the *G. lemaneiformis* culture area (Fig. 2a). In this study, *V. alginolyticus* was verified to be the dominant bacterium on rotten algae infected by ‘Baotou’ disease (Fig. 1a). In contrast, *V. alginolyticus* was not detected on the healthy samples of H1 and H3, and accounted just 0.56% of the total community on H2 (Fig. 1a). Besides, the *V. alginolyticus* accounted 7.5% of the total microbial community in the seawater near ‘Baotou’ samples, which was significantly higher than that in the heathy samples (0.18%, Fig 2b). The results of bacterial communities showed that *V. alginolyticus* might have a close relationship with the outbreak of ‘Baotou’ disease.

ZB10 exhibited the highest pathogenicity in the 10 strains of bacteria which could cause the rotting of *G. lemaneiformis.* Moreover, the symptom caused by the ZB10 was the rotting of the main trunk of the algae, that was consistent with the ‘Baotou’ disease occurring in farms. Sun et al (2011). Have reported in a detailed study that two strains of pathogenic bacteria, *Thalassospira* sp. and *Vibrio parahaemolyticus*, can cause tip bleaching of *G. lemaneiformis.* The alga with ‘Baotou’ disease has healthy tips, while its trunk is the main rotting part (Fig 3b), that is different from the characteristics reported by Sun et al. (2011). Liu et al. (2019) also isolated and screened three strains of *Agarivorans albus*, *Aquimarina latercula* and *Brachybacterium* sp. that can cause bleaching of *G. lemaneiformis*, but the morphological characteristics of the pathogenic alga were not reported in detail in that study.

ZB10 was identified as *V. alginolyticus* by morphological and molecular identification. Its morphology was the same as that reported by Zhang et al (31). The results of the microbial community experiments and the symptoms caused by the *V. alginolyticus* both indicated that *V. alginolyticus* was the pathogen of the ‘Baotou’ disease. Besides, a large number of *V. alginolyticus* cells were found to be attached to the decaying thalli surface of *G. lemaneiformis* after 72 h of co-culturing and infection with the *V. alginolyticus*. In addition, *V. alginolyticus* can be easily re-isolated from the decayed algal thalli infected in the lab. Furthermore, a large number of rod-shaped and spherical bacterial cells identified as *V. alginolyticus* by morphological identification, were found to be attached to the ‘Baotou’ disease infected thalli collected from the farm (Fig 4 f). Based on the Koch’s postulates and the above results (bacterial communities, the same decaying symptoms, *V. alginolyticus* can be easily re-isolated from the decayed algal thalli infected in the lab), the *V. alginolyticus* was identified as the pathogen of ‘Baotou’ disease.

Interestingly, *V. alginolyticus* is the pathogen to ‘Baotou’ diseased *G. lemaneiformis* in this study, but a beneficial bacterium to *S. japonica* reported reduces the bleaching disease risk of *S. japonica* (18). *G. lemaneiformis* and *S. japonica* are often co-cultured in China due to their growth periods overlapped for about 3 months. In the past ten years, the breeding scale of *G. lemaneiformis* and *S. japonica* has continued to expand rapidly. The total production of *G. lemaneiformis* and *S. japonica* in China have exceeded 2.1 million tons in 2022(8). These two species of seaweeds are often plagued with bleaching diseases during their high-density commercial cultivation in recent years (12,32). *V. alginolyticus* is the pathogen to ‘Baotou’ diseased *G. lemaneiformis* in this study, but a beneficial bacterium to *S. japonica*. The study seems to remind us that *V. alginolyticus* is closely involved in the competition or co-existence between *G. lemaneiformis* and *S. japonica* co-cultured in China. *V. alginolyticus* appears in large quantities in the culture farms of *G. lemaneiformis* and *S. japonica*, and is pathogenic to *G. lemaneiformis* but beneficial to *S. japonica*. Clarifying the ecological and biological mechanisms of this phenomenon might be very interesting and meaningful for ensuring the large-scale production of seaweeds and understanding the ecological balance between different seaweeds. Besides, *V. alginolyticus* is also pathogenic to other mariculture organisms such as *Haliotis diversicolor*, *Ruditapes decussatus* and *Litopenaeus vannamei* (33,34,35). Cai et al. (2006) reported an astonishing 90% mortality rate of *H. diversicolor* postlarvae attributed to *V. alginolyticus*. Interestingly, both *G. lemaneiformis* and *S. japonica* are the main feed for *H. diversicolor* in China. Gomez-Leon et al. (2005) found that within a period of 30 d, the mortality rate of *R. decussatus* soared to 60% due to the presence of *V. alginolyticus*. Ngo et al. (2020) reported that the survival rate of *L. vannamei* infected with *V. alginolyticus* was significantly reduced to only 50%, in comparison to that of the unaffected control group. Considering the rich ecological role of *V. alginolyticus* and its close relationship with aquatic economic species, more research should be devoted to it to further clarify its biological and ecological characteristics.

Collectively, *V. alginolyticus* was the dominant bacterium on *G. lemaneiformis* with ‘Baotou’ disease. In contrast, there were very few *V. alginolyticus* on the healthy samples. *V. alginolyticus* could cause the rotten symptoms of *G. lemaneiformis* in the lab, and the symptoms it caused were consistent with those of ‘Baotou’ disease. Besides, *V. alginolyticus* can be easily re-isolated from the rotten algal thalli infected in the lab. A large number of *V. alginolyticus* cells were found to be attached to the algal thalli with ‘Baotou’ disease and the rotten thalli acquired by the lab *V. alginolyticus* infection treatment. *V. alginolyticus* strictly conforms to the four criteria of Koch’s postulates regarding pathogens, and was proven to be the pathogen causing ‘Baotou’ disease in cultivated *G. lemaneiformis* (36,37).

## Materials and Methods

### Source of experimental seaweed

*G. lemaneiformis* used for the experiment was collected from the Ailun Bay farm in Rongcheng City, Shandong Province, China (37°10′ N, 122°34′ E) between June to September 2023. Experiments were carried out immediately after they were collected.

### Collection of the bacterial community samples associated with the G. lemaneiformis

The healthy and rotten thalli (5 g) were putted in 50ml tubes respectively and then incubated in the liquid nitrogen immediately after they were collected (3 replicates for each group). Then the healthy (H) and rotten (R) thalli were used for studying the bacteria communities attached to the *G. lemaneiformis*. Seawater about 0.5 meters away from the healthy and rotten *G. lemaneiformis* thalli was collected respectively. Then 1 liter of seawater was filtered by 0.2μm filter membranes respectively to obtain bacterial community samples in the seawater around the healthy (HS) and rotten (RS) *G. lemaneiformis*. The filter membranes were putted in 5ml tubes respectively and incubated in the liquid nitrogen immediately after the bacteria were collected (3 replicates for each group).

### Analysis of the bacterial community structure

Genomic DNA of bacteria was extracted using the Power Soil DNA Isolation Kit (DNeasy) according to manufacturer’s instructions. The 16S rRNA full sequence regions was amplified by PCR. Purified products were pooled in equimolar and DNA library was constructed using the SMRT bell prep kit 3.0 (Pacifc Biosciences, CA, USA) according to PacBio’s instructions. Purified SMRT bell libraries were sequenced on the Pacbio Sequel IIe System (Pacifc Biosciences, CA, USA) by Majorbio Bio-Pharm Technology Co. Ltd. (Shanghai, China). High-fidelity (HiFi) reads were obtained from the subreads, generated using circular consensus sequencing via SMRT Link v11.0. 485518 sequences were denoised using the DADA2 (26) plugin in Quantitative Insights Into Microbial Ecology (QIIME2 v.2020.2) (27). After denoising, 217602 sequences (ranging from 10105 to 27654 per sample) were obtained, and finally these sequences were clustered into 1443 Amplicon Sequence Variants (ASVs). Taxonomic assignment of ASVs was performed using the Vsearch consensus taxonomy classifier implemented in Qiime2 and the SILVA 16S rRNA database (v138). Stacked bar plot conducted by R (v3.3.1) was used to identify the most abundance bacterial communities on species levels, the top 30 species in terms of abundance in the samples were shown.

### Isolation of pathogenic bacteria

The rotten seaweed tissue (0.2 g) was homogenized in 3 ml of sterile seawater for 3 minutes at room temperature (23 °C) then was centrifuged at 800g for 5 min. After centrifugation, 0.1 mL of supernatant was diluted by 10, 100, 1000, and 10000 times, and 0.1 ml of the different dilutions were spread onto Tryptic Soy Broth (TSB) solid culture medium. After 48 hours of cultivation (23 °C), single bacterial colony was isolated by TSB plate streaking cultivation, and this process was repeated three times to obtain pure bacterium.

### Pathogenicity assessment

Acquired single bacterial colonies were picked from the solid culture medium and transferred into 50 ml of sterile TSB liquid culture medium. After 24 hours of cultivation at 23°C in a shaker (shake rate of 110 rpm), the cultures were centrifuged at 12,000g for 10 minutes. Sediment was collected and diluted in 500 ml of sterile seawater to obtain a diluted bacterial suspension. Then, healthy algal thalli (weighting 5 g) were incubated in the bacterial suspension (500 ml) for 8 hours at 21°C in the dark. After incubation, the seaweed (5g) was cultivated in different breeding ponds containing 200 L seawater (32 psu, changed every day), at a temperature 21°C, light 60μmol m^-2^s^-1^ at 14L:10D photoperiod. According to the method of Pang et al. (2024), the number of rotten points on each algal thallus was recorded at 0, 24, 48 and 72h. 2 replicates were set for each single bacterial colony, along with 2 control groups consisting of 500 ml of sterile seawater.

After the first time of preliminary pathogenicity assessment for each bacterial colony, those colonies exhibiting pathogenicity were subjected to further infection validation test. Ten replicates were set for each bacterial colony, along with 10 control groups. The number of rotten points on each algal thallus was recorded at 0, 24, 48 and 72 h according to the method of Pang et al (12).

### 16S rRNA Sequencing and phylogenetic analysis of isolated bacteria

Total DNA was extracted from 1ml bacterial solution using Ezup Column Bacteria Genomic DNA Purification Kit, following the kit protocol. Then, 16s rRNA was amplified using the method that Gu et al described (28). Successful PCR products were purified and sequenced by a commercial company (Sangon, China). The Blast algorithm at the National Center for Biotechnology Information (http://www.ncbi.nlm.gov/blast/) was used to compare the nucleotide sequences. Relevant sequences were obtained from GenBank for phylogenetic analysis. A maximum likelihood (ML) phylogenetic tree was constructed with the MEGA 11 program (29). The reliability of the branching was tested by bootstrap re-sampling (1000 pseudo-replicates).

### Observation of bacterial morphology on seaweed

Rotten seaweeds (3 replicates for each type) were collected from the farm and lab pathogenicity assessment experiment, keeping they in their original state, and then proceeded with scanning electron microscopy observation immediately. The processing procedure followed the method described by Pang et al (12). Then, samples were observed by scanning electron microscopy (Hitachi S-3400).

### Observation of pathogenic bacterial morphology

Transmission electron microscope (TEM) was used to observe the cell morphology of bacteria. The sample preparation method was modified from Ma et al (18). Bacteria suspension (OD_600_=1.3, 20ml) was centrifugation at 3,000 g for 10 minutes and resuspended in 2.5% (wt/vol) glutaraldehyde for 10 minutes. Then, 10μl of suspension was moved to a carbon-coated copper grid for 10 minutes. Finally, the sample was observed by transmission electron microscope Hitachi-HT7700.

## Statistical analysis

Statistical analyses were performed using IBM PASW Statistics 18. One-way ANOVA and the least significant difference (LSD) at *p*<0.05 was used to test the significant differences.

## Author contributions

F.W.: Performed the experiments, Investigation, Methodology, Statistical analysis. N.Y.: Investigation, discussion of the results. T.P.: Conceptualization, Methodology, Writing – review & editing, Supervision, Funding acquisition. Q.G.: Statistical analysis, Writing - review & editing. Y.S.: Investigation, discussion of the results. J.L.: Writing - review, Funding acquisition.

## Data availability

Data are available on request to the corresponding author.

## References

1. Yun JH, Archer SD, Price NN. 2022. Valorization of waste materials from seaweed industry: An industry survey based biorefinery approach. Rev Aquacult 15:1020–1027. 10.1111/raq.12748

2. Khan N, Sudhakar K, Mamat R. 2024. Macroalgae farming for sustainable future: Navigating opportunities and driving innovation. Heliyon 10:e2820 8. 10.1016/j.heliyon.2024.e28208

3. Zhou X. 2022. The State of World Fisheries and Aquaculture 2022. Towards Blue Transformation. Rome, FAO 44–45. 10.4060/cc0461en

4. Pérez-Estrada CJ, León-Tejera H, Serviere-Zaragoza E. 2012. Cyanobacteria and macroalgae from an arid environment mangrove on the east coast of the Baja California Peninsula. botm 55:187-196. 10.1515/bot-2012-0501

5. Wen X, Peng C, Zhou H, Lin Z, Lin G, Chen S, Li P. 2006. Nutritional Composition and Assessment of *Gracilaria lemaneiformis* Bory. J Integr Plant Biol 48(9):1047−1053. 10.1111/j.1744-7909.2006.00333.x

6. Chen X, Tang Y, Sun X, Zhang X, Xu N. 2022. Comparative transcriptome analysis reveals the promoting effects of IAA on biomass production and branching of *Gracilariopsis lemaneiformis*. Aquaculture 548. 10.1016/j.aquaculture.2021.737678

7. Xie X, He Z, Hu X, Wang Q, Yang Y. 2023. The composition, function and assembly mechanism of epiphytic microbial communities on *Gracilariopsis lemaneiformis*. J Exp Mar Biol Ecol 564. 10.1016/j.jembe.2023.151909

8. Liu X Z. 2023. China Fishery Statistical Yearbook. China Agriculture Press.

9. Jiang H, Zou D, Lou W, Chen W, Yang Y. 2019. Growth and photosynthesis by *Gracilariopsis lemaneiformis* (Gracilariales, Rhodophyta) in response to different stocking densities along Nan’ao Island coastal waters. Aquaculture 501:279–284. 10.1016/j.aquaculture.2018.11.047

10. Liu X, Chen Y, Zhong M, Chen W, Lin Q, Du H. 2019. Isolation and pathogenicity identification of bacterial pathogens in bleached disease and their physiological effects on the red macroalga *Gracilaria lemaneiformis*. Aquat Bot 153:1–7. 10.1016/j.aquabot.2018.11.002

11. Sun X, He Y, Xu N, Xia Y, Liu Z. 2011. Isolation and identification of two strains of pathogenic bacteria and their effects on the volatile metabolites of *Gracilariopsis lemaneiformis* (Rhodophyta). J Appl Phycol 24:277–284. 10.1007/s10811-011-9677-0

12. Pang T, Wang F, Yang X, Liu J, Xu H, Guo Q, Liu L. 2024. A contagious disease caused serious damage to *Gracilariopsis lemaneiformis* cultivation in China. J Appl Phycol 36:2005–2011 (2024). 10.1007/s10811-024-03246-6

13. Pang T, Zhang M, Lu L, Liu J. 2022. Hypo-osmotic soaking shows potential as a selective method of removing *Ceramium filicula*, the dominant epiphyte causing serious damage to *Gracilariopsis lemaneiformis* cultivation in China. Aquaculture 561:738642. 10.1016/j.aquaculture.2022.738642

14. Yang Y, Chai Z, Wang Q, Chen W, He Z, Jiang S. 2015. Cultivation of seaweed *Gracilaria* in Chinese coastal waters and its contribution to environmental improvements. Algal Res 9:236–244. 10.1016/j.algal.2015.03.017

15. Ward GM, Faisan JP, Cottier-Cook EJ, Gachon C, Hurtado AQ, Lim PE, Matoju I, Msuya FE, Bass D, Brodie J. 2019. A review of reported seaweed diseases and pests in aquaculture in Asia. J World Aquacult Soc 51:815–828. 10.1111/jwas.12649

16. Ding M. 1991. The effects of the environmental factors on Laminaria disease caused by alginic acid decomposing bacteria. Acata Oceanol Sin 11(1):123–130.

17. Wang G, Shuai L, Li Y, Lin W, Zhao X, Duan D. 2007. Phylogenetic analysis of epiphytic marine bacteria on Hole-Rotten diseased sporophytes of Laminaria japonica. J Appl Phycol 20:403–409. 10.1007/s10811-007-9274-4

18. Ma M, Zhuang Y, Chang L, Xiao L, Lin Q, Qiu Q, Chen D, Egan S, Wang G. 2024. Naturally occurring beneficial bacteria *Vibrio alginolyticus* X-2 protects seaweed from bleaching disease. mbio 14:1-19. 10.1128/mbio.00065-23

19. Largo DB, Fukami K, Nishijima T. 1995. Occasional pathogenic bacteria promoting ice-ice disease in the carrageenan-producing red algae *Kappaphycus alvarezii* and *Eucheuma denticulatum* (Solieriaceae, Gigartinales, Rhodophyta). J Appl Phycol 7:545–554. 10.1007/BF00003941

20. Largo DB, Fukami K, Nishijima T. 1999. Time-dependent attachment mechanism of bacterial pathogen during ice-ice infection in *Kappaphycus alvarezii* (Gigartinales, Rhodophyta). J Appl Phycol 11: 129–136. 10.1023/A:1008081513451

21. Syafitri E, Prayitno SB, Ma’ruf WF, Radjasa OK. 2017. Genetic diversity of the causative agent of ice-ice disease of the seaweed *Kappaphycus alvarezii* from Karimunjawa island, Indonesia. Earth Environ Sci 55:012044. 10.1088/1755-1315/55/1/012044

22. Yang R, Liu Q, He Y, Tao Z, Xu M, Luo Q, Chen J, Chen H. 2020. Isolation and identification of *Vibrio mediterranei* 117-T6 as a pathogen associated with yellow spot disease of *Pyropia* (Bangiales, Rhodophyta). Aquaculture 526:735372. 10.1016/j.aquaculture.2020.735372

23. Ding H, Ma J. 2005. Simultaneous infection by red rot and chytrid diseases in *Porphyra yezoensis* Ueda. J Appl Phycol 17:51–56. 10.1007/s10811-005-5523-6

24. Lavilla-Pitogo. 1992. Agar-digesting bacteria associated with ‘rotten thallus syndrome’ of *Gracilaria* sp. Aquaculture 102:1–7. 10.1016/0044-8486(92)90283-q

25. Martinez JN, Padilla PIP. 2016. Isolation and characterization of agar-digesting *Vibrio* species from the rotten thallus of *Gracilariopsis heteroclada* Zhang et Xia. Mar Environ Res 119:156–160. 10.1016/j.marenvres.2016.05.023

26. Amir A, McDonald D, Navas-Molina JAKopylova E, Morton JT, Zech Xu Z, Kightley EP, Thompson LR, Hyde ER, Gonzalez A, Knight R. 2017. De blur rapidly resolves single-nucleotide community sequence patterns. mSystems 2:10.1128/msystems.00191-16. https://doi.org/10.1128/msystems.00191-16

27. Segata N, Izard J, Waldron L, Gevers D, Miropolsky L, Garrett WS, Huttenhower C. 2011 Metagenomic biomarker discovery and explanation. Genome Biol 12:R60 (2011). 10.1186/gb-2011-12-6-r60

28. Gu Y, Zheng R, Sun C, Wu S. 2022. Isolation, Identification and Characterization of Two Kinds of Deep-Sea Bacterial Lipopeptides Against Foodborne Pathogens. Front Microbiol 13:792755. 10.3389/fmicb.2022.792755

29. Kumar V, Zozaya-Valdes E, Kjelleberg S, Thomas T, Egan S. 2016. Multiple opportunistic pathogens can cause a bleaching disease in the red seaweed *Delisea pulchra*. Environ Microbiol 18:3962–3975. 10.1111/1462-2920.13403

30. Zhao Y, Zhou N, Ren J, Liu W, Zhou C, Chen X, Zhao J, Cao J, Yang J, Han J, Liu H. 2023. Comprehensive insights into the metabolism characteristics of small RNA Qrr4 in *Vibrio alginolyticus*. Appl Microbiol Biotechnol 107:1887–1902. 10.1007/s00253-023-12435-1

31. Zhang J, Liu K, Gong X, Zhang N, Zeng Y, Ren W, Huang A, Long H, Xie Z. 2023. Transcriptome analysis of the hepatopancreas from the *Litopenaeus vannamei* infected with different flagellum types of *Vibrio alginolyticus* strains. Front Cell Infect Microbiol 13:1265917. 10.3389/fcimb.2023.1265917

32. Zhang R, Chang L, Xiao L, Zhang X, Han Q, Li N, Egan S, Wang G. 2020. Diversity of the epiphytic bacterial communities associated with commercially cultivated healthy and diseased *Saccharina japonica* during the harvest season. J Appl Phycol 32:2071–2080. 10.1007/s10811-019-02025-y

33. Cai J, Han H, Song Z, Li C, Zhou J. 2006. Isolation and characterization of pathogenic *Vibrio alginolyticus* from diseased postlarval abalone, Haliotis diversicolor supertexta (Lischke). Aqua Res 37:1222–1226. 10.1111/j.1365-2109.2006.01552.x

34. Ngo HV, Huang HT, Lee PT, Liao ZH, Chen HY, Nan FH. 2020. Effects of Phyllanthus amarus extract on nonspecific immune responses, growth, and resistance to *Vibrio alginolyticus* in white shrimp *Litopenaeus vannamei*. Fish Shellfish Immunol 107:1–8. 10.1016/j.fsi.2020.09.016

35. Gomez-Leon J, Villamil L, Lemos ML, Novoa B, Figueras A. 2005. Isolation of *Vibrio alginolyticus* and *Vibrio splendidus* from aquacultured carpet shell clam (*Ruditapes decussatus*) larvae associated with mass mortalities. Appl Environ Microbiol 71:98–104. 10.1128/AEM.71.1.98-104.2005

36. Koch R. Uber bakteriologische forschung verhandlung des X internationalen medichinischen congresses, Berlin, 1890, 1, 35. August Hirschwald, Berlin. (In German.) Xth International Congress of Medicine, Berlin. 1891.

37. Byrd AL, Segre JA. 2016. Adapting Koch’s postulates. Science 351, 224–226. 10.1126/science.aad6753

